# Temporal clustering of sleep spindles and coupling with slow oscillations in the consolidation and generalization of motor memory

**DOI:** 10.1101/2024.07.26.605262

**Authors:** Adrien Conessa, Damien Léger, Arnaud Boutin

**Affiliations:** Université Paris-Saclay, INRIA, CIAMS, Orsay, France, 91405; VIFASOM, Université Paris Cité, Paris, France, 75004; Centre du Sommeil et de la Vigilance, APHP, Hôtel-Dieu, Paris, France, 75004

**Keywords:** Sleep, spindles, slow oscillations, memory consolidation, sensorimotor restriction, immobilization, infraslow

## Abstract

Sleep benefits memory consolidation through periodic sleep spindle activity and associated memory reactivations. Recent evidence highlights that iterative spindle onsets follow two periodic rhythms: an infraslow periodicity (∼0.02 Hz) and a mesoscale periodicity rhythm (∼0.2-0.3 Hz). Consequently, spindles tend to cluster in “trains” on a low-frequency time scale every 50 seconds during which spindles iterate every 3 to 4 seconds. Such temporal organization of spindles in trains is considered a critical sleep mechanism for the timed and repeated reactivation of memories. Additionally, current trends indicate that a timely phase-locking between slow oscillations (SO) and spindles promotes learning-related synaptic plasticity. In this study, we explored how spindle clustering and coupling with SO contribute to motor memory consolidation by inducing a local reduction in synaptic efficacy over sensorimotor cortical regions through upper-limb immobilization after motor sequence learning. We also evaluated memory generalization using two transfer tests designed to assess the ability to transfer or generalize the newly acquired skill to another one (new sequence) or another effector (inter-limb transfer). Our results reveal that the temporal cluster-based organization of spindles is independent of daytime sensorimotor experience, while distinct overnight behavioral outcomes were elicited. Interestingly, immobilization induced a phase shift in the SO-spindle coupling for spindles grouped in trains, but not when isolated outside trains. In addition, the proportion of grouped spindles relative to isolated spindles was positively associated with skill consolidation and negatively correlated with skill generalization following sensorimotor restriction. These results suggest that spindle trains may promote skill-specific strengthening of motor memories, while isolated spindles may instead create memory-instability conditions that facilitate skill generalization.

## Introduction

Behind every learned motor skill lies a period of practice and rehearsal (1, 2). However, although the repeated practice of a motor skill is crucial for its initial acquisition, developing an effective movement representation is not only a result of practice (3). The newly formed memory continues to be processed “offline” during waking and sleeping hours following practice. This offline period offers a privileged time window for memory consolidation, which relates to the process whereby newly acquired and relatively labile memories are transformed into enhanced and more stable memory traces (4, 5). Following its initial acquisition, the memory trace is thought to be dynamically maintained during wakefulness and actively reprocessed during a subsequent sleep period, allowing its strengthening and thus influencing its generalizability (3, 6–8). A daytime nap or a night of sleep has long been shown to play a crucial role in the strengthening and transformation of motor memories during consolidation (9–12).

Another important aspect of motor learning involves the ability to generalize the acquired skill to another one or another effector, such as inter-limb transfer. This fundamental principle underlies our capacity to perform unpracticed motor skills or learn new ones more easily (7). While the role of sleep in the skill-specific strengthening of motor memories during consolidation has been widely studied, evidence also showed that it can promote their generalization (7, 13, 14). Indeed, recent theoretical evidence suggests that learning contexts creating conditions of memory instability may be critical for the generalization of motor skills (7, 14). Instability is thought to make a memory vulnerable to interference and even forgetting, following its initial formation or reactivation, with the loss of detailed knowledge reducing memory specificity and enabling the development of generalizable knowledge (7). Hence, it remains to be determined whether sleep consolidation mechanisms may offer optimal conditions for motor memory strengthening and memory-instability conditions supporting the creation of generalized knowledge (4, 7, 15).

Growing evidence suggests that brief bursts of sigma-like thalamocortical oscillations, known as sleep spindles, during non-rapid eye movement (NREM) sleep play a critical role in the offline covert reactivation of motor memories during consolidation, resulting in overnight/nap memory improvements (9, 11, 14, 16–19). Sleep spindle activity is an electrophysiological hallmark of NREM-stage 2 (NREM2) sleep involving short (0.3-2 sec) synchronous bursts of waxing and waning 11-16 Hz oscillations (8, 14, 18). Boutin and colleagues (2020) (8) recently developed a framework for motor memory consolidation that outlines the essential contribution of the clustering and hierarchical rhythmicity of spindle activity during this sleep-dependent process. More specifically, sleep spindles tend to cluster in ‘trains’ on a low-frequency time scale of approximately 50 seconds (∼0.02 Hz infraslow rhythm) during NREM sleep, interspersed with relative spindle-free periods where sporadic isolated spindles may occur. This infraslow periodicity has been considered a crucial neurophysiological regulatory mechanism providing transient temporal windows of optimal neural conditions for the reactivation of memory traces through spindle activity (20–22). In addition to the periodic clustering of spindles, recent findings unveiled a recurring pattern where spindles tend to reoccur approximately every 3-4 seconds during trains (∼0.2-0.3 Hz mesoscale rhythm). In theory, such inter-spindle intervals during trains represent periods of refractoriness that are assumed to be crucial for the timely and organized segregation of spindle-induced memory reactivations, thus regulating the cyclic interference-free reprocessing of memory traces for efficient memory consolidation (8, 18, 23, 24). However, recent evidence suggests that spindle trains during NREM2 sleep play a more significant role in motor memory consolidation compared to those occurring during NREM3 sleep (18).

Accompanying this multi-scale rhythmicity of spindle activity, it has been proposed that cross-frequency coupling and hierarchical nesting of NREM sleep rhythms may be critical for memory consolidation. Current trends postulate that slow oscillations (SO; 0.5-1.25 Hz) confer a temporal window for spindles to occur in their excitable up-states and that a timely phase-locking facilitates the induction of persistent synaptic plastic changes (25–32) (see (33) for a review). At the synaptic level, post-learning neuroplasticity has further been shown to be mediated by inter-hemispheric sleep regulations over localized sensorimotor regions (19, 34, 35), suggesting that local regulations of both SOs and spindles reflect learning-dependent brain plasticity and skill consolidation during sleep. Indeed, slow-wave activity (0.5-4 Hz) is now established as a marker of homeostatically regulated sleep pressure and synaptic downscaling during NREM sleep (36). In an elegant study, Huber and colleagues (2006) (35) transiently promoted local synaptic depression in sensorimotor regions using a short-term upper-limb immobilization paradigm. Results revealed that the immobilization procedure induced local homeostatic changes (i.e., reduced spectral power mainly in the SO frequency band) in the contralateral affected-limb sensorimotor cortex that were associated with deteriorations in motor performance. Moreover, Debarnot and colleagues (2021) (37) revealed that sleep spindle activity is also locally affected by immobilization-induced synaptic depression. Thus, upper-limb immobilization has proven to be an effective paradigm for investigating the causal link between daytime sensorimotor experience and sleep-dependent memory consolidation, as it induces cortical synaptic changes that modulate sleep characteristics (35, 37–39). Recent studies further demonstrated that a fine-tuned SO-spindle coupling depends on prior sensorimotor experience and structural brain integrity (28, 29, 40). Hence, considering that SOs and spindles are involved in synaptic consolidation, the potential contribution of spindles’ clustering and the precision of the SO-spindle coupling in the strengthening and generalization of motor skills during consolidation remains to be explored.

Using a short-term upper-limb immobilization procedure, combined with the recording of behavioral and electroencephalographic (EEG) night sleep measures, the aim of the present study was twofold: (i) to determine the effects of daytime sensorimotor experience on local sleep spindle expression (i.e., clustering and rhythmicity) and the integrity of the cross-frequency coupling between SOs and spindles, and (ii) to evaluate their functional contribution in the consolidation and generalization of motor skills. Given that upper-limb immobilization disrupts the balance of inter-hemispheric inhibition (39), a process closely tied to motor learning and generalization (41), it can be hypothesized that transient upper-limb immobilization would impact the consolidation and generalization of memory traces. Adopting a time-based clustering approach, we hypothesized that the rhythmic occurrence of spindles in trains would confer favorable conditions for efficient reprocessing and consolidation of the memory trace, supported by an effective plasticity-dependent SO-spindle coupling compared to spindles occurring in isolation. Isolated spindles are instead expected to impair skill-specific motor memory strengthening through ineffective, non-recurring memory reactivations (18) or impaired SO-spindle coupling, thus rendering the memory trace unstable and prone to generalization.

## Results

Thirty right-handed participants were equally divided into a left upper-limb immobilization group (IMMO) or a control group without immobilization (CTRL). A 13-hour immobilization procedure was administered immediately after the practice of a finger motor-sequence task (using the non-dominant left hand) to transiently induce a local reduction in synaptic efficacy within the contralateral sensorimotor cortical regions (35). Sleep EEG recordings were collected the night before (acclimatization night) and after motor sequence learning (experimental night). Behavioral performance was assessed during motor learning, as well as during test blocks before and after the experimental night (Figure 1). The ability to transfer or generalize the newly acquired skill to another one (new sequence) or another effector (i.e., inter-limb transfer) was also assessed after the experimental night. Analyses were conducted on the temporal cluster-based organization of sleep spindles, SO-spindle coupling, and their relationship with behavioral performance. See Materials and methods for further details.

**Figure 1.**
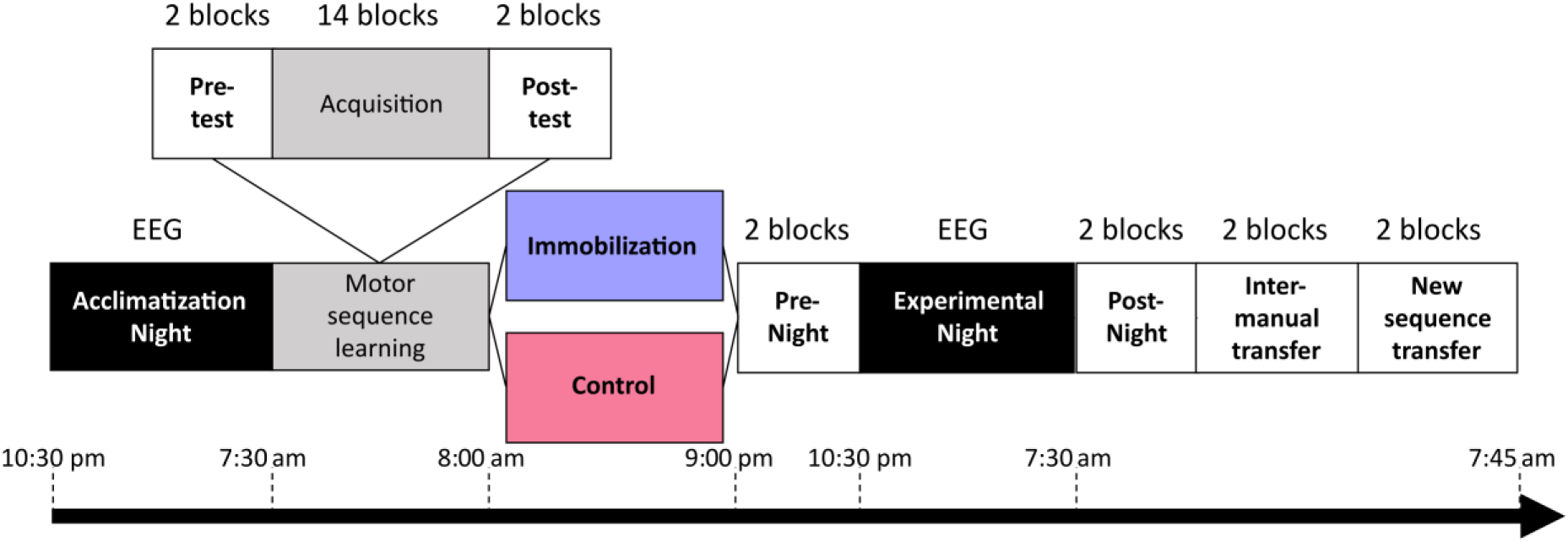
Experimental design. Sleep EEG recordings of 30 participants were acquired during two consecutive nights. Between the two nights, in the morning, participants were trained on an 8-element finger movement sequence. Immediately after training, one-half of the participants were left-hand immobilized (n = 15, IMMO group) for a period of 13 hours, while the other half were not (n = 15, CTRL group). Following the immobilization or control procedure, all participants were tested on the trained motor sequence during pre-night and post-night tests. They were then evaluated on two transfer tests: an inter-manual transfer test (same motor sequence performed with the right-hand fingers), and a new motor-sequence test (unpracticed motor sequence performed with the left-hand fingers). EEG: Electroencephalography.

### The immobilization procedure only reduced the use of the immobilized upper limb

To control for the correct immobilization of the left upper limb in the IMMO group, a mixed ANOVA was performed on the mean velocity score computed from the accelerometer with the between-subject factor CONDITION (IMMO, CTRL) and the within-subject factor LATERALITY (Left, Right). The analysis revealed significant main effects for the factor CONDITION (F(1,28) = 21.0, *p* < 0.001, ɳ²p = 0.43), and LATERALITY (F(1,28) = 164, *p* < 0.001, ɳ²p = 0.85), as well as a significant CONDITION x LATERALITY interaction (F(1,28) = 111, *p* < 0.001, ɳ²p = 0.80). Holm post-hoc tests revealed that the velocity score of the left limb was lower in the IMMO group than in the CTRL group (*p* < 0.001), as well as compared to the right limb of the CTRL group (*p* < 0.001) and IMMO group (*p* < 0.001), which did not differ from each other (*p* = 0.75). These findings thus guarantee compliance with the immobilization procedure in the IMMO group, whose participants used their left upper limb significantly less than their right upper limb, and less than both upper limbs in the CTRL group.

### Sensorimotor restriction only impaired inter-manual skill transfer

Figure 2 shows the mean response times measured for each test block. Analyses regarding the number of accurately typed sequences are available in Supplementary Information (Figure S1). A mixed ANOVA using a CONDITION (IMMO, CTRL) x BLOCK (pre-test, post-test) factorial design, with repeated measures on the BLOCK factor, was first performed to ensure that the two groups did not differ in terms of performance before and after the acquisition phase. The analysis revealed a significant effect of the factor BLOCK (F(1, 28) = 102, *p* < 0.001, ɳ²p = 0.79), indicating performance improvements during training for both groups. The analysis failed to detect a significant effect of the factor CONDITION (F(1, 28) = 0.93, *p* = 0.34, ɳ²p = 0.03), nor a BLOCK x CONDITION interaction (F(1, 28) = 0.44, *p* = 0.51, ɳ²p = 0.02), suggesting that performance before the immobilization procedure did not differ between the IMMO and CTRL groups.

**Figure 2.**
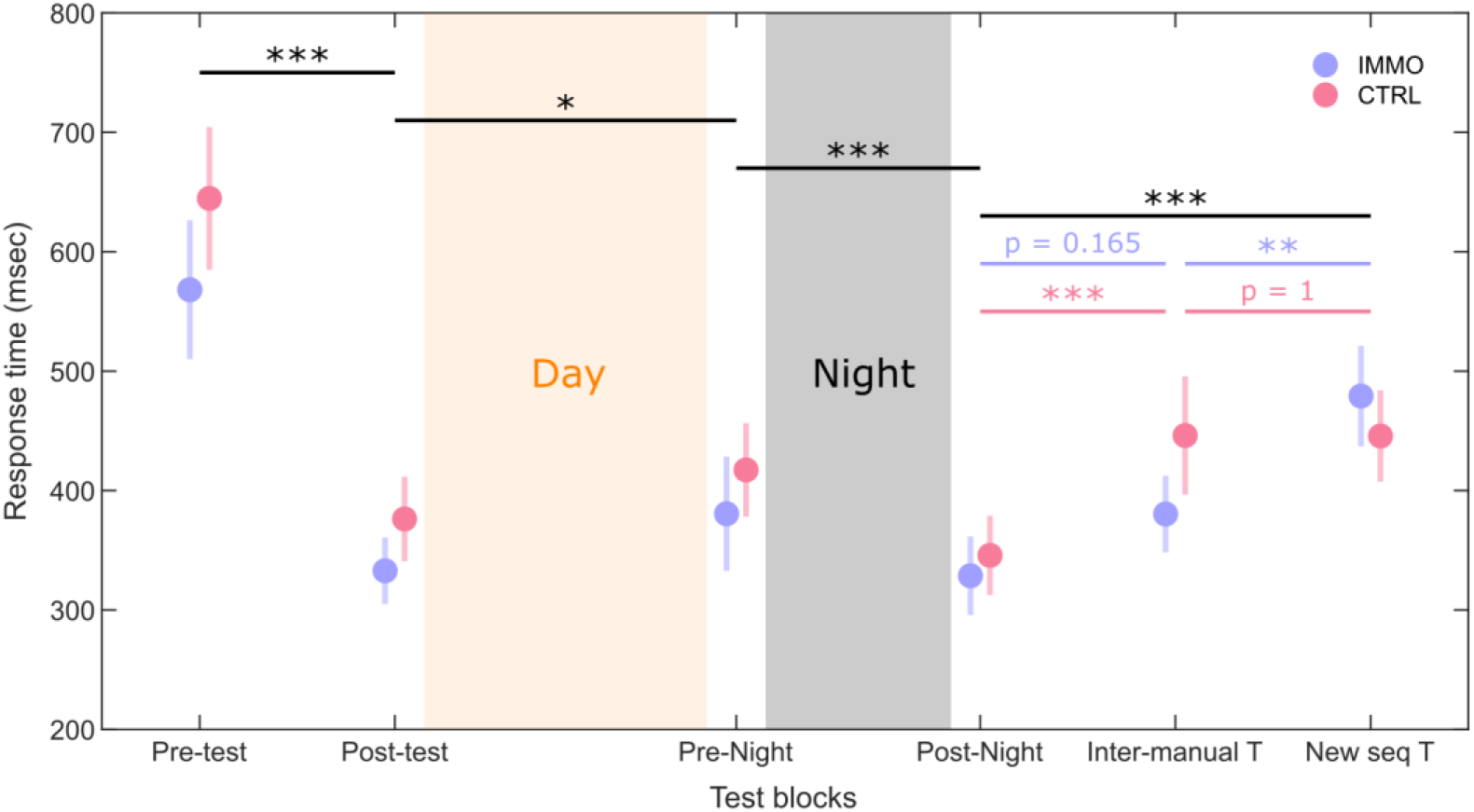
Sensorimotor restriction only impaired inter-manual skill transfer. Mean response time for each test block in the IMMO group (purple) and CTRL group (pink). The colored circles represent the group average, and the bar represents the standard error of the mean (n = 15 for each group and each test block, with the exception of the transfer test blocks with n = 13 for the IMMO group). The stars represent p-values associated with mixed ANOVA tests. * p < 0.05, ** p < 0.01, *** p < 0.001. Inter-manual T: Inter-manual transfer; New seq T: new-sequence transfer.

To compare changes in performance over the 13-hour retention interval, a mixed ANOVA was performed using a CONDITION (IMMO, CTRL) x BLOCK (post-test, pre-night) factorial design, with repeated measures on the factor BLOCK. The analysis revealed a significant main effect of the factor BLOCK (F(1, 28) = 7.30, *p* = 0.012, ɳ²p = 0.21), suggesting a deterioration in performance during the day for both groups. The analysis failed to detect a main effect of the factor CONDITION (F(1, 28) = 0.60, *p* = 0.44, ɳ²p = 0.02) and a CONDITION x BLOCK interaction (F(1, 28) = 0.04, *p* = 0. 84, ɳ²p < 0.01).

To compare changes in performance after the experimental night, a mixed ANOVA was performed using a CONDITION (IMMO, CTRL) x BLOCK (pre-night, post-night) factorial design, with repeated measures on the factor BLOCK. The analysis revealed a significant main effect of the factor BLOCK (F(1, 28) = 29.4, *p* < 0.001, ɳ²p = 0.51) but failed to detect a main effect of the factor CONDITION (F(1, 28) = 0.25, *p* = 0.62, ɳ²p < 0.01) nor a CONDITION x BLOCK interaction (F(1, 28) = 0.74, *p* = 0.40, ɳ²p = 0.03). Thus, both groups improved after the night, but overnight skill consolidation were not significantly different.

To compare skill generalizability between groups, a mixed ANOVA was performed using a CONDITION (IMMO, CTRL) x BLOCK (post-night, inter-manual transfer, new-sequence transfer) factorial design, with repeated measures on the factor BLOCK. Two participants in the IMMO group were removed from this analysis as they did not perform the new-sequence transfer test. The analysis revealed a significant effect of the factor BLOCK (F(2, 52) = 30.9, *p* < 0.001, ɳ²p = 0.54) and a significant CONDITION x BLOCK interaction (F(2, 52) = 4.64, *p* = 0.014, ɳ²p = 0.15) without detecting a significant main effect of the factor CONDITION (F(1, 26) = 0.19, *p* = 0.67, ɳ²p < 0.01). Holm’s post-hoc comparisons revealed significant performance decreases from the post-night test to the inter-manual transfer test in the CTRL group, but not in the IMMO group. Also, impaired skill generalization toward a new motor sequence was revealed for both groups, as expressed by significant performance decreases from the post-night test to the new-sequence transfer test. Finally, the IMMO group showed a significant difference in performance between the inter-manual transfer test and the new-sequence transfer, in contrast to the CTRL group.

### Sensorimotor restriction induced local reduction of synaptic efficacy

To assess whether the short-term sensorimotor restriction led to a reduction in synaptic efficacy, a Student-t permutation test for independent samples (1000 permutations) was performed to obtain the statistical topographic map of the spectral power difference in the slow oscillation frequency band between the IMMO and CTRL groups (Figure 3). The analysis revealed significant decreases in SO power in the IMMO group than in the CTRL group on multiple electrodes, which are primarily located over sensorimotor cortical regions contralateral to the immobilized limb, after correction for multiple comparisons using the Benjamini-Hochberg procedure (42) to control the false discovery rate (Ntest = 63). Hence, the present result shows a general decrease in the spectral power of slow oscillations during the first 20 minutes of NREM2 sleep following the 13 hours of upper-limb immobilization. We also performed supplementary analyses on the C4 derivation, showing that the difference in the SO power between the two groups fades when considering the whole night (Figure S2).

**Figure 3.**
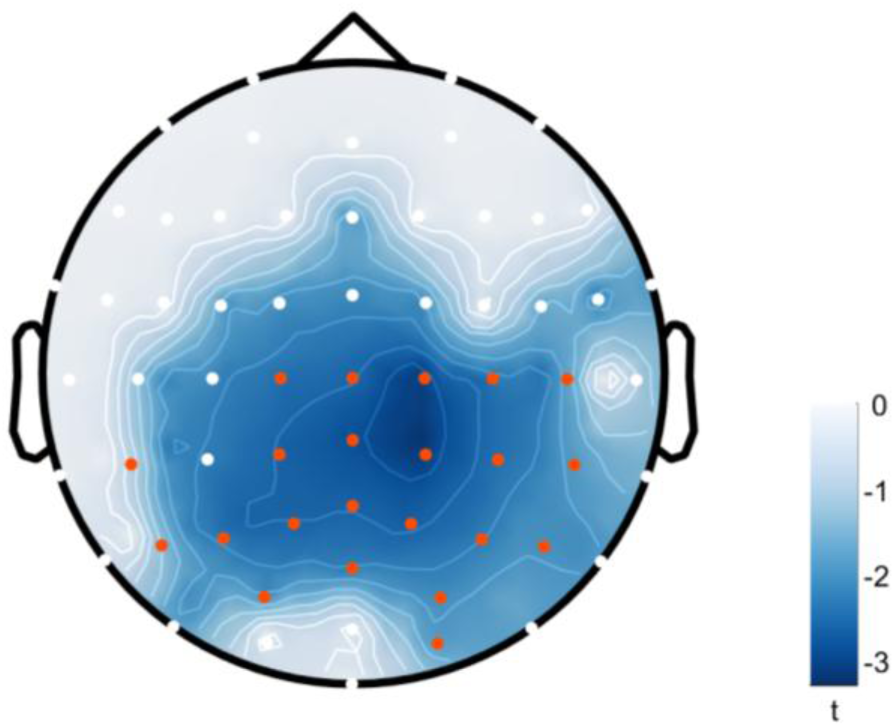
Sensorimotor restriction of the left upper limb decreased the spectral power of the slow oscillation frequency band over the contralateral right sensorimotor cortex. Topographical statistical map of the difference in SO (0.5–1.25 Hz) spectral power between the IMMO and CTRL groups during the first 20 minutes of NREM2 sleep (IMMO-CTRL contrast). The color bar represents the t-test values. Negative t values (blue) represent lower spectral power for the IMMO group (n = 15) compared with the CTRL group (n = 15). White areas represent the non-significant differences between groups after the Student-t permutation test (1000 permutations). The red dots highlight the electrodes with statistical differences after correction for multiple comparisons using the Benjamini-Hochberg procedure to control the false discovery rate (Ntest = 63).

### Sensorimotor restriction disrupted the beneficial effect of grouped spindles on skill consolidation and favored skill generalization conveyed by isolated spindles

To assess the specific role of grouped and isolated sleep spindles in the skill consolidation process, correlation analyses were performed between the proportion of grouped over isolated spindles during the first 20 minutes of NREM2 sleep, extracted at scalp derivation C4, and the magnitude of skill consolidation (percentage of performance changes from the pre-night test to the post-night test). One participant in the CTRL group did not show grouped spindles in the first 20 minutes of NREM2 sleep and was therefore excluded from the correlation analyses. A significant positive relationship was observed between the proportion of grouped spindles during the first 20 minutes of NREM2 sleep and the magnitude of skill consolidation in the CTRL group only (r = 0.58, *p* = 0.031) but not in the IMMO group (r = −0.10, *p* = 0.72) (Figure 4A). We also aimed to compare the coefficient correlations between groups by conducting Fisher’s r to z transform and computing the statistical significance of the observed z-test statistic from the differences of the transformed z scores. No significant difference was found between the IMMO and CTRL groups (Z = 1.83, p = 0.07). However, it is noteworthy that large sample sizes (N = 66) are needed to detect large-sized differences of Pearson correlation coefficient (Δr ∼ 0.5) (43). Considering r values of .10, .30, and .50 as the thresholds for small, medium, and large effect sizes, respectively (43), our correlation analyses are in favor of greater involvement of grouped than isolated spindles in the memory consolidation process for the CTRL group. No significant relationship was found when the proportion of grouped spindles was computed over the whole night in the CTRL (r = −0.23, *p* = 0.41) and IMMO groups (r = 0.03, *p* = 0.92).

**Figure 4.**
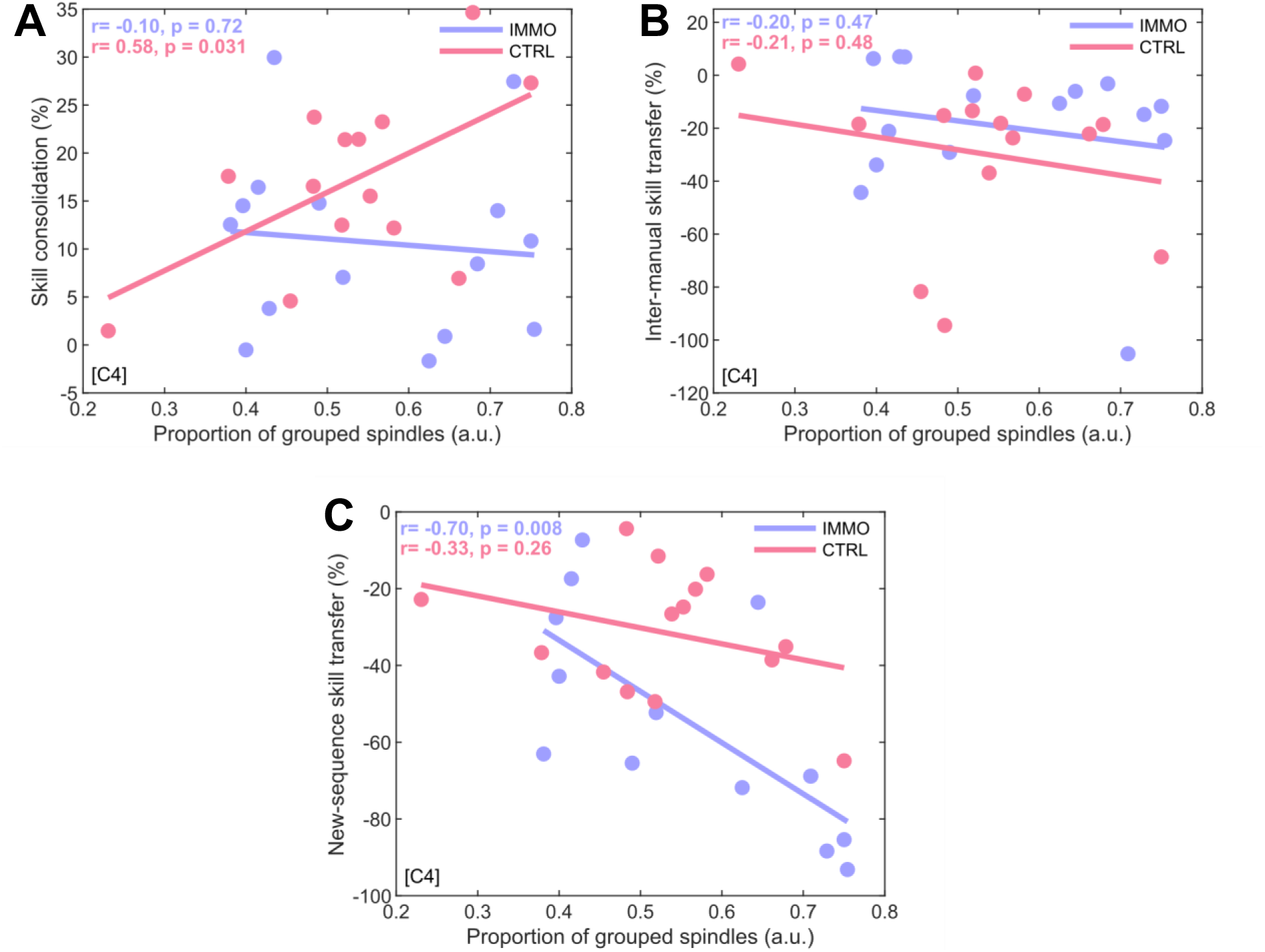
Sensorimotor restriction disrupted the beneficial effect of grouped spindles on skill consolidation and favored skill generalization conveyed by isolated spindles. **(A)** Relationship between the proportion of grouped spindles over isolated spindles (a.u.), extracted at scalp derivation C4, and the magnitude of overnight skill consolidation (in %) on the learned motor sequence in the IMMO (n = 15) and CTRL (n = 14) groups. **(B)** Relationship between the proportion of grouped spindles over isolated spindles (a.u.), extracted at scalp derivation C4, and the magnitude of inter-manual skill transfer (in %) in the IMMO (n = 15) and CTRL (n = 14) groups. **(C)** Relationship between the proportion of grouped spindles over isolated spindles (a.u.), extracted at scalp derivation C4, and the magnitude of new-sequence skill transfer (in %) in the IMMO (n = 13) and CTRL (n = 14) groups. Individual data (colored circles) and trend lines are provided, along with Pearson’s r and associated p-values.

We also assessed the role of grouped spindles in the ability to generalize the acquired skill to another one or another effector (inter-limb transfer). To that end, we separately correlated the proportion of grouped over isolated spindles during the first 20 minutes of NREM2 sleep, extracted at scalp derivation C4, with the magnitude of i) inter-manual skill transfer (percentage of performance changes from the post-night test to the inter-manual transfer test) and ii) new-sequence skill transfer (percentage of performance changes from the post-night test to the new-sequence transfer test). As mentioned previously, two participants of the IMMO group did not undergo the new-sequence transfer test and were therefore excluded from this analysis. No significant relationship was found between the proportion of grouped spindles and the magnitude of inter-manual skill transfer in the IMMO group (r = −0.20, *p* = 0.47) and in the CTRL group (r = −0.21, *p* = 0.48) (Figure 4B). Similarly, no significant relationship was found when the proportion of grouped spindles was computed over the whole night in the CTRL (r = 0.23, *p* = 0.41) and IMMO groups (r = −0.27, *p* = 0.33). Interestingly, though, the analysis revealed a significant negative relationship between the proportion of grouped spindles and the magnitude of new-sequence skill transfer in the IMMO group only (r = −0.70, *p* = 0.008) but not in the CTRL group (r = −0.33, *p* = 0.26) (Figure 4C). Hence, we also aimed to compare the coefficient correlations between groups by conducting Fisher’s r to z transform. No significant difference was found between the IMMO and CTRL groups (Z = 1.20, p = 0.23). Considering r values of .10, .30, and .50 as the thresholds for small, medium, and large effect sizes, respectively (43), our correlation analyses are in favor of greater involvement of isolated than grouped spindles in the memory generalization process for the IMMO group; a higher proportion of isolated spindles enhances the ability to transfer or generalize the newly acquired skill to another one (new sequence). No significant relationship was found when considering the whole night, in the CTRL (r = 0.04, *p* = 0.89) and IMMO groups (r = −0.32, *p* = 0.29). Additional analyses regarding the effect of sensorimotor restriction on sleep and spindle characteristics are provided in Supplementary Information (see Tables S1 and S2 for details on the sleep architecture and spindle characteristics of the experimental and acclimatization nights, respectively).

### Sensorimotor restriction affected the SO-spindle coupling

Phase-amplitude coupling analyses were performed to assess whether the upper limb’s transient immobilization influences the coupling between slow oscillations and sleep spindles. These analyses were first performed on the SO-spindle concomitant events detected during the first 20 minutes of NREM2 sleep, at scalp derivation C4 (Figure 5A and B). Participants without spindles coupled to slow oscillations during this period were removed from the current analysis (2 in the IMMO group, 4 in the CTRL group). The Rayleigh and the Watson-Williams tests were applied to test the non-uniformity of the preferred coupling phases and to compare them between the IMMO and CTRL groups. Analyses indicated a non-uniform coupling in both IMMO (θ = 0 rad, Rayleigh Z = 9.54, *p* < 0.001) and CTRL groups (θ = −0.83 rad, Rayleigh Z = 7.97, *p* < 0.001). A significant difference in the preferred coupling phase was observed (F(1,22) = 11.9, *p* = 0.002) during the first 20 minutes of NREM2 sleep, showing a delay in the coupling of slow oscillations and sleep spindles for participants in the IMMO group compared to their CTRL group counterparts.

**Figure 5.**
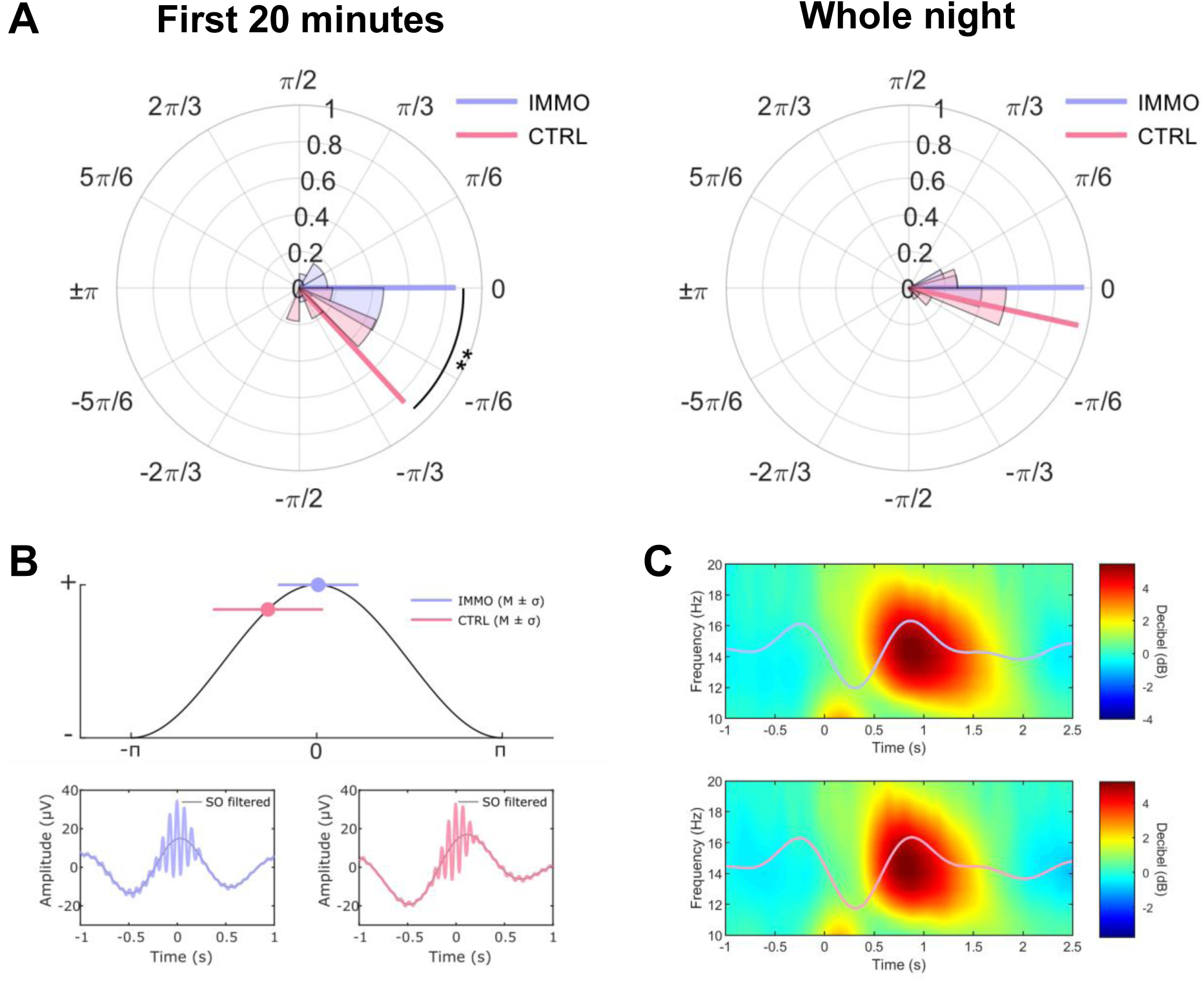
Sensorimotor restriction affected SO-spindle coupling during the first part of the night. **(A)** Preferred phase in radians (and mean length of the resultant vector) of slow oscillations at the peak power of sleep spindles, detected at scalp derivation C4, for the IMMO (purple) and CTRL (pink) groups during NREM2 sleep when considering the first 20 minutes (left) and the whole night (right). Probability density plots of phases (histograms) are also provided. **(B)** Top. Representation of the mean preferred phase and the circular standard deviation of the SO-spindle coupling for the IMMO and CTRL groups on a simulated slow oscillation. Bottom. Peak-locked sleep spindle group average across all concomitant events during the first 20 minutes of NREM2 sleep, with the bandpass-filtered SO trace (in gray) superimposed to highlight the SO-spindle coupling in both the IMMO (left) and CTRL (right) groups. **(C)** Time-frequency maps (in dB) in the 10-20 Hz frequency range of all concomitant events detected at scalp derivation C4 during all NREM2 epochs over the whole night, using epoch windows ranging from −1 to +2.5 sec around slow oscillation onset. Time-frequency maps were displayed with the average bandpass-filtered SO trace superimposed to illustrate the high synchronization between sleep spindle power and slow oscillation phase in the IMMO (top) and CTRL (bottom) groups. The stars represent p-values associated with the Watson-William test. ** p < 0.01

The limited number of participants showing SO-spindle coupling with both types of spindles (grouped and isolated) during the first 20 minutes of NREM2 sleep does not allow us to perform such clustering analyses. Therefore, separate analyses of the SO-spindle coupling for grouped and isolated spindles were only performed on all artifact-free NREM2 epochs over the entire sleep recording (whole night).

First, for all NREM2 sleep spindles, a non-uniform SO-spindle coupling was found for participants in both the IMMO (θ = 0 rad, Rayleigh Z = 13.8, *p* < 0.001) and CTRL groups (θ = −0.22 rad, Rayleigh Z = 13.6, *p* < 0.001). No significant group difference in the preferred phase was observed (F(1,28) = 3.77, *p* = 0.062), showing the absence of phase shift in the SO-spindle coupling between the IMMO and CTRL groups when considering all NREM2 sleep spindles of the whole night (Figure 5A and C).

Second, for all NREM2 grouped spindles, a non-uniform SO-spindle coupling was found for participants in both the IMMO (θ = 0.24 rad, Rayleigh Z = 8.90, *p* < 0.001) and CTRL groups (θ = −0.23 rad, Rayleigh Z = 12.2, *p* < 0.001). A significant group difference in the preferred phase was observed (F(1,28) = 4.25, *p* = 0.049), suggesting that a phase shift in the SO-spindle coupling between the IMMO and CTRL groups is present when considering only grouped spindles (Figure 6).

**Figure 6.**
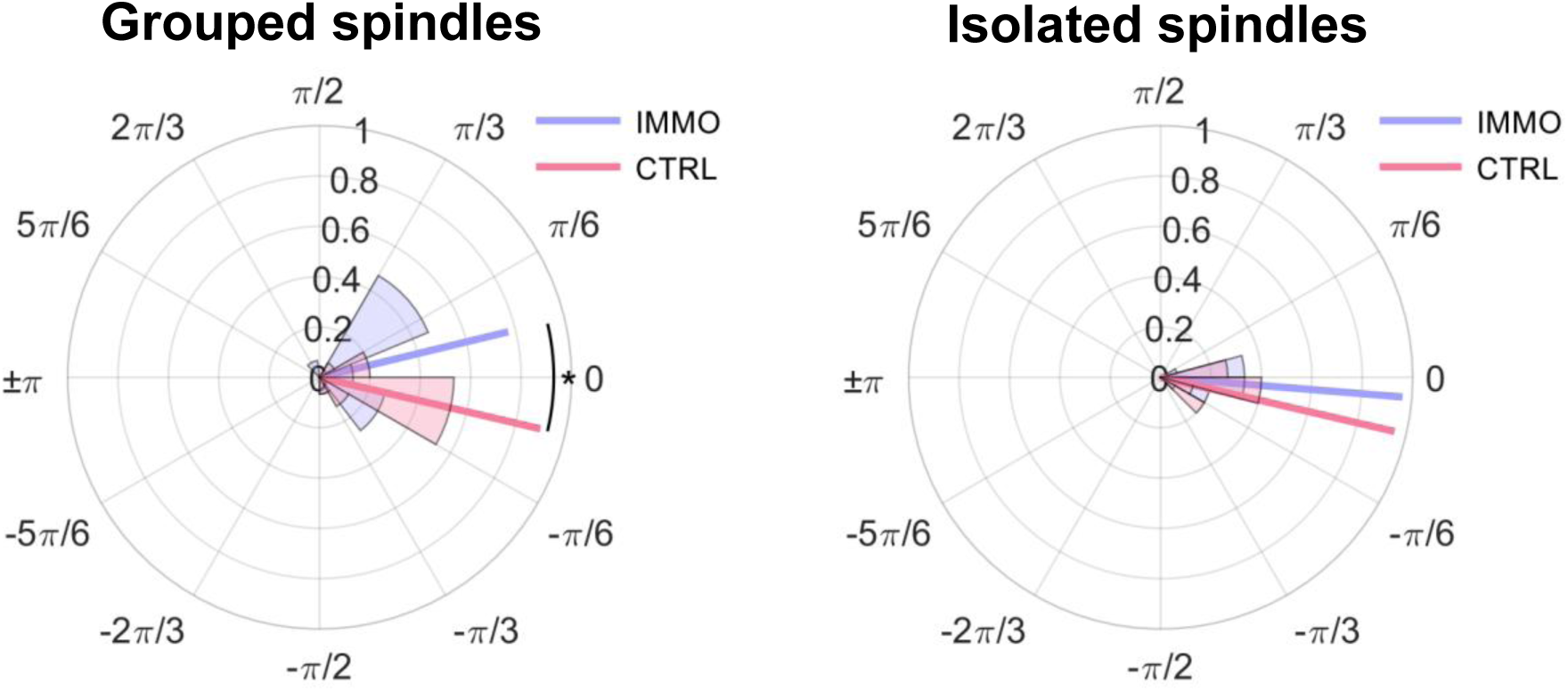
A phase-shift in the SO-spindle coupling was induced only for grouped spindles. Preferred phase in radians (and mean length of the resultant vector) of slow oscillations at the peak power of sleep spindles, detected at scalp derivation C4, for the IMMO (purple) and CTRL (pink) groups during NREM2 sleep of the whole experimental night when considering only grouped spindles (left) or isolated spindles (right). Probability density plots of phases (histograms) are also provided. The star represents the p-value associated with the Watson-William test. * p < 0.05.

Third, for all NREM2 isolated spindles, a non-uniform SO-spindle coupling was found for participants in both the IMMO (θ = −0.08 rad, Rayleigh Z = 13.9, *p* < 0.001) and CTRL groups (θ = −0.23 rad, Rayleigh Z = 13.6, *p* < 0.001). Interestingly, though, no significant group difference in the preferred phase was observed (F(1,28) = 1.76, *p* = 0.20), showing the absence of phase shift in the SO-spindle coupling between the IMMO and CTRL groups when considering only isolated spindles (Figure 6).

These results suggest that synaptic changes induced by transient immobilization delayed the onset of the power peak of spindles occurring in trains (grouped spindles) but not when occurring in isolation. No difference was observed between the IMMO and CTRL groups on the acclimatization night (Figure S3). Similar analyses were also carried out on other electrodes (Figure S4). This SO-spindle shift between the IMMO and CTRL groups was found for a few electrodes, mainly located over the right sensorimotor cortex when considering only grouped spindles, suggesting a local effect of the immobilization on the SO-spindle coupling.

### Multi-scale fluctuations of spindle-band power were not affected by sensorimotor restriction

The temporal organization of sleep spindles is governed by an infraslow (∼0.02 Hz) and a mesoscale rhythm (∼0.2–0.3 Hz). The infraslow and mesoscale spectral profiles were computed for all artifact-free NREM2 sleep epochs during the whole experimental night (Figure 7). An independent Student t-test was then performed between the IMMO and CTRL groups to compare the effect of the immobilization on the peak power frequency of the infralow periodicity. The analyses failed to reveal a significant difference (t(28) = 1.05, *p* = 0.30, d = 0.39), suggesting that the periodicity of spindle trains (infraslow rhythm) did not differ between the IMMO (P = 0.017 ± 0.004 Hz) and CTRL groups (P = 0.019 ± 0.005 Hz).

**Figure 7.**
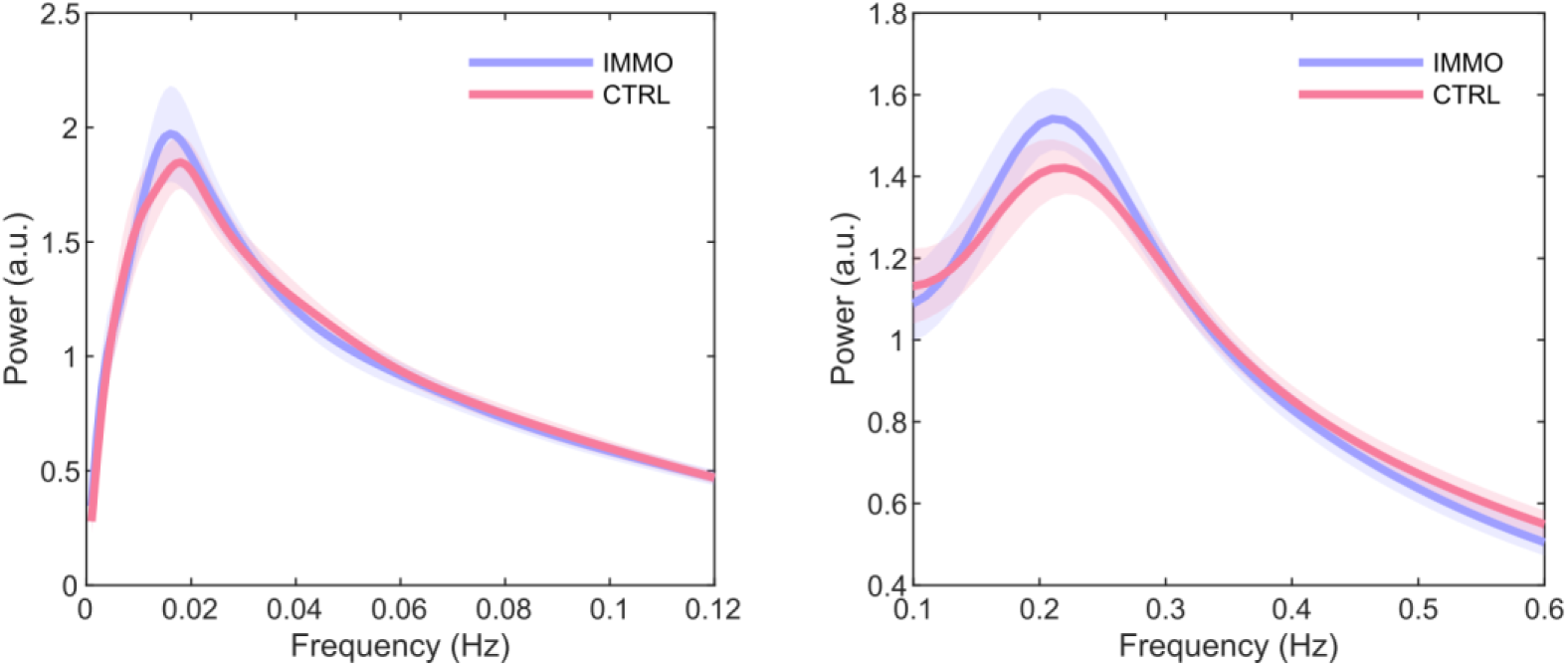
Multi-scale fluctuations of spindle-band power were not affected by sensorimotor restriction. Grand average of the spectral profile at the scalp derivation C4, ranging from 0.001 to 0.12 Hz (left panel) and 0.1 to 0.6 Hz (right panel) frequency range of the sleep spindle frequency band (11– 16 Hz) power fluctuation during NREM2 sleep episodes across the whole night. The peaks around 0.02 Hz and 0.2 Hz for both the IMMO (purple) and CTRL (pink) groups highlight the multi-scale periodicity of spindle-band power at an infraslow and mesoscale rhythms, respectively. The lighter areas surrounding the mean represent the standard error of the means.

An independent Student t-test was also performed between the IMMO and CTRL groups to compare the effect of the immobilization on the peak power frequency of the mesoscale periodicity. The analyses failed to reveal a significant difference (t(28) < 0.01, *p* = 1, d < 0.01), suggesting that the periodicity of spindles within trains (mesoscale rhythm) did not differ between the IMMO (P = 0.21 ± 0.03 Hz) and CTRL groups (P = 0.21 ± 0.04 Hz). The present results accord with previous findings, confirming the hierarchical multi-scale periodicity in sleep spindle activity with spindle-band power fluctuations at frequencies close to 0.02 Hz and 0.2–0.3 Hz. Here, the immobilization procedure did not affect the double periodicity of the spindle-band power during NREM2 sleep.

Similar analyses were performed for the acclimatization night (Figure S5) and failed to detect significant differences, highlighting the stability of the multi-scale sleep spindle periodicity before and after the immobilization procedure.

## Discussion

In the current study, we examined (i) the effects of daytime sensorimotor experience on local sleep spindle expression (i.e., clustering and rhythmicity) and the integrity of the cross-frequency coupling between SOs and spindles, and (ii) their functional contribution in the consolidation and generalization of motor skills. On the one hand, our findings corroborate previous work (34, 35) by showing significant local decreases in the SO spectral power during the first 20 minutes of sleep in the IMMO group (IMMO-CTRL contrast), mainly over the affected sensorimotor cortex (i.e., contralateral to the immobilized limb), reflecting the induction of local synaptic depression following immobilization (35). At the behavioral level, this primarily manifested as enhanced skill generalization to a different effector. Despite the immobilization-induced synaptic depression, our results did not reveal any significant difference between the IMMO and CTRL groups regarding spindle clustering and rhythmicity: sleep spindles tend to cluster on a low-frequency time scale of about 50 seconds (∼0.02 Hz rhythm), during which spindles iterate every 3-4 seconds (∼0.2-0.3 Hz rhythm), irrespective of daytime sensorimotor experience. On the other hand, alterations of the coupling between SOs and spindles during NREM2 sleep have been shown to occur locally over the affected sensorimotor cortex mainly during the first 20 minutes of sleep but not when considering the whole night. Interestingly, immobilization induced a phase shift in the SO-spindle coupling for grouped spindles but not for isolated spindles, supporting the hypothesis of their functional differences in memory consolidation. More specifically, our results accord with recent theories assuming that sleep spindles occurring in trains (grouped spindles) play a more critical role than isolated ones in motor skill consolidation. In contrast, isolated spindles may impair skill consolidation by providing conditions of memory instability that support the creation of generalized knowledge.

At the behavioral level, our findings revealed that both groups deteriorated performance over the day. Early learning results in a degradation of performance throughout the day, potentially due to the “early-boost” phenomenon (44), or a reduction in the signal-to-noise ratio of the newly acquired memory trace (45). However, immobilization of the upper limb did not significantly worsen performance deterioration during the day. This result may be surprising at first glance, but the deleterious effect of immobilization has mainly been shown on mental rotation (46, 47) and pointing tasks (35, 48), in contrast to motor sequence tasks (49). For the latter, the primary motor cortex seems to have a functional role in the early stage of consolidation but not in the more delayed stages (a few hours after learning) (50). Therefore, the delayed synaptic depression induced by immobilization over the contralateral sensorimotor cortex may not have been early enough to disrupt the early phase of motor skill consolidation. Likewise, transient limb immobilization did not affect the later stages of motor skill consolidation, expressed behaviorally by the magnitude of (overnight) skill retention performance. We conjecture that sensorimotor restriction may have induced the reactivation of additional unaffected brain regions during sleep, acting as a compensatory consolidation mechanism to mitigate the deleterious effects of local synaptic depression. Indeed, in accordance with previous work (49, 51, 52), our results indicated that the IMMO group did not perform significantly worse on the inter-manual transfer test than on the post-night test, unlike the CTRL group. The potential overuse of the non-immobilized limb in the IMMO group may have reduced the inter-hemispheric inhibition produced by the affected hemisphere (39) and, in turn, increased the excitability of the contralateral hemisphere, leading to enhanced inter-manual skill transfer.

In line with previous work (34, 35), a significant decrease in the SO spectral power was found over the sensorimotor cortex of the immobilized limb during the first 20 minutes of NREM2 sleep in the IMMO group (IMMO-CTRL contrast). Interestingly, this early-night difference in SO power between the two groups fades when considering the whole night (Figure S2). Hence, SO activity may be viewed as a sensitive marker of synaptic efficacy, modulated by daytime sensorimotor experience (34, 35, 53). In our study, the decrease in SO activity suggests that the immobilization procedure likely induced local synaptic depression during the following night, reducing synaptic efficacy in the sensorimotor cortex of the immobilized limb, as previously described (35, 37–39).

Despite this immobilization-induced synaptic depression, our results did not reveal any significant difference between the IMMO and CTRL groups regarding spindle clustering and rhythmicity. As previously shown, current results confirmed the clustering and temporal organization of spindle activity during NREM2 sleep, irrespective of daytime sensorimotor experience: spindles tend to cluster on a low-frequency time scale of about 50 seconds (∼0.02 Hz rhythm), during which spindles iterate every 3-4 seconds (∼0.2-0.3 Hz rhythm) (8, 14, 18, 20, 54). Thus, the temporal organization of spindles in trains appears to be an inherent clocking sleep mechanism independent of prior sensorimotor experience, regulated in a learning-independent manner and coordinated with autonomic nervous system-dependent parameters (8, 14, 18, 20, 22, 55). Yet, in line with recent findings of Boutin and colleagues (8, 14, 18, 19), an in-depth analysis of spindle clustering revealed a significant positive relationship between the proportion of grouped spindles over the sensorimotor cortex of the left limb during the first 20 minutes of sleep and the overnight skill consolidation for the CTRL group but not for the IMMO group. The findings indicate that local synaptic depression caused by immobilization may have impaired the efficacy of grouped spindles, thereby disrupting the beneficial effects of repeated memory trace reactivations. Interestingly, correlation analyses were significant only when considering the proportion of grouped spindles at the beginning of the night. From a theoretical perspective, because the relevant memory trace was acquired in the morning, we may propose that sleep spindles selectively reactivate memory traces in a (pre-)defined temporal order that depends on the degree of synaptic plasticity induced by daytime sensorimotor experience. In line with the synaptic homeostasis hypothesis (36) that emphasizes the role of sleep in regulating synaptic weight in the brain, supposed to be maximal at the beginning of sleep and then progressively returning to baseline, brain regions with higher synaptic efficacy following learning should be processed first during subsequent sleep.

In contrast, however, we demonstrate a significant negative relationship between the proportion of grouped spindles and the magnitude of new-sequence skill transfer in the IMMO group but not in the CTRL group. This suggests that spindles occurring in isolation during NREM2 sleep may facilitate skill generalizability toward a new (related) motor skill. This finding aligns with recent studies supporting the hypothesis that sleep spindles might also be related to forgetting (18, 56–58). More specifically, Boutin and colleagues (2024) (18) recently suggested that spindles occurring in trains can strengthen memory representations through their timed reactivations, while in contrast, spindles occurring in isolation may instead activate sleep mechanisms promoting memory-instability conditions leading to the clearance or decreased accessibility of the memory content. Furthermore, since the infraslow rhythm is also related to arousability, segmenting sleep into periods of high arousability (for environmental alertness) and low arousability (for memory consolidation) (20), it is tempting to suggest that sporadic reactivations triggered by isolated spindles during periods of heightened alertness might result in inefficient reactivations, which could ultimately destabilize and weaken the memory trace (7). At the same time, recent theoretical evidence suggests that memory instability may be critical for motor skill generalization (7). Thus, immobilization-induced synaptic depression may have led to impaired reprocessing of the memory trace during sleep through local alterations related to spindle clustering, giving way to its weakening by isolated spindles and, consequently, its generalization. Altogether, our results accord with recent theories assuming that sleep spindles occurring in trains (grouped spindles) play a more critical role than isolated ones in motor skill consolidation by offering optimal conditions for motor memory strengthening, while isolated spindles may instead promote memory-instability conditions supporting the creation of generalized knowledge (4, 7).

Finally, alterations of the coupling between SOs and spindles during NREM2 sleep have been shown to occur locally over the affected sensorimotor cortex mainly during the first 20 minutes of sleep, suggesting that the synaptic depression induced by transient immobilization delayed the onset of the power peak of sleep spindles. Interestingly, this delay is only observed for grouped spindles and not for isolated spindles, and it persists throughout the night. Solano et al. (2022) (59) recently demonstrated that the proportion of grouped spindles coupled to slow oscillations increased locally in the sensorimotor cortex of the trained effector, thus suggesting that the SO-spindle coupling relies on previous sensorimotor experience. Hence, it is likely that altered local synaptic efficacy at the cortical level disrupted the SO-spindle coupling mostly during spindle trains. However, the origin of this altered phase shift is uncertain, as immobilization-induced cortical neuroplasticity is assumed to occur early in the night (35). In a previous study, Helfrich and colleagues (2018) (29) reported a phase shift in the SO-spindle coupling for older adults, compared to young adults, associated with a memory consolidation deficit. This phase shift was linked to a structural alteration of gray matter in the medial prefrontal cortex. In addition, it was described that the structural integrity of the thalamocortical fascicle influences sleep spindle density (60). For example, daytime learning and sensorimotor experience can rapidly induce white matter changes (61), increasing sleep spindle production during the subsequent night (60). Based on the observation that sensorimotor restriction also induces white matter structural changes (62), the transient alteration of white matter bundles may be a potential cause of the SO-spindle phase shift. This latter hypothesis remains speculative, and further evaluations of the effects of sensorimotor restriction on sleep spindle expression are needed. Nevertheless, the specificity of this effect on grouped spindles highlights the potential existence of distinct neurobiological mechanisms underlying and modulating the generation of grouped and isolated spindles, supporting the hypothesis of their functional differences in memory consolidation (63).

Our findings confirm the multiscale periodicity of sleep spindle activity regarding spindle clustering and rhythmicity. We further revealed that this temporal cluster-based organization of sleep spindles is not dependent on daytime sensorimotor experience. Furthermore, by experimentally inducing a phase shift in the coupling between SO and sleep spindles occurring in trains, but not in isolation, our results support the theoretical assumption that grouped and isolated spindles may have distinct functional roles in memory consolidation. While sleep spindles occurring in trains may be involved in motor skill consolidation by offering optimal conditions for memory strengthening, isolated spindles may instead be involved in the weakening or even forgetting of memories by creating conditions of memory instability, which may in turn favor the development of generalized knowledge.

## Data availability

The data that support the results of this study are available from the corresponding author upon reasonable request and under a formal data-sharing agreement

## Code availability

Sleep EEG data were processed using the MATLAB R2021b software from The MathWorks (Natick, MA) and the open-source Brainstorm software (64). The codes for the detection and clustering of sleep spindles are available at the following GitHub repositories: https://github.com/arnaudboutin/Spindle-detection and https://github.com/arnaudboutin/Spindle-clustering. The codes for slow oscillations and infraslow-mesoscale analyses are available at https://github.com/arnaudboutin/Spindle-SO-package. The codes used to perform other analyses are available from the corresponding author upon request.

## Acknowledgments

We thank the medical and technician staff of the Hôtel-Dieu sleep Center for providing facilities and for their assistance in organizing the study. This work was partly supported by a research grant from the “Société Française de Physiothérapie” to financially compensate participants for their involvement in this study.

## Materials and methods

### Participants

Thirty healthy volunteers (16 females, mean age: 25.4 ± 4 years) were recruited by local advertisements and were randomly and equally divided into two groups: an immobilization group (IMMO; n = 15, 7 females, mean age: 25.9 ± 4 years) and a control group without immobilization (CTRL; n = 15, 9 females, mean age: 25.0 ± 4 years). All participants met the following inclusion criteria: aged between 18 and 35 years, right-handed (Edinburgh Handedness Inventory (65)), medication-free, without history of mental illness, epilepsy, or head injury with loss of consciousness, sleep or neurologic disorders, and no recent upper extremity injuries. The experimental protocol was approved by the “Comité de Protection des Personnes Sud-Ouest et Outre-Mer III” (ID-RCB: 2020-A01465-34) and conformed to relevant guidelines and regulations. All participants gave written informed consent before inclusion. Participants were asked to maintain a regular sleep-wake cycle and to refrain from all caffeine- and alcohol-containing beverages 24 hours prior to the experimentation.

### Experimental design

Participants sat on a chair at a distance of 50 cm in front of a computer screen. The motor task consisted of performing as quickly and accurately as possible an 8-element finger movement sequence by pressing the appropriate response keys on a standard French AZERTY keyboard using their left, non-dominant hand fingers. The sequence to be performed (B-C-N-V-C-B-V-N, where C corresponds to the little finger and N to the index finger) was explicitly taught to the participant before training.

Test and practice blocks consisted of repeating the 8-element sequence for 30 seconds. Each block began with the presentation of a green cross in the center of the screen accompanied by a brief 50-ms tone. In case of occasional errors, participants were asked “not to correct errors and to continue the task from the beginning of the sequence” (see (14) for a similar procedure). At the end of each block, the color of the green imperative stimulus turned red, and participants were then required to look at the fixation cross during the 30-s rest period. This experimental protocol was designed to control the duration of practice during the test and practice blocks. Stimuli presentation and response registration were controlled using the MATLAB R2019b software from The MathWorks (Natick, MA) and the Psychophysics Toolbox extensions (66).

Each participant completed two visits at the Sleep and Vigilance Center of the Hotel-Dieu Hospital (Figure 1). The first visit served as a polysomnographic (PSG) screening and acclimatization night, whereas the second visit consisted of the experimental night following motor sequence learning and the daytime immobilization or non-immobilization control procedure. Assignment to the immobilization or control group was randomized across participants. The first visit started at 9:00 pm; the experimental design was explained and all the forms were provided. After the participants had prepared for the night, the EEG equipment was set up and they were invited to sleep. The sleep EEG recording started approximately at 10:30 pm. The EEG cap was removed the next morning at 6:30 am, just after awakening. The experimental procedure began at 7:30 am for all participants to minimize the possible impact of circadian and homeostatic factors on individual performance and give them time to shower and have breakfast. The experimental procedure for the first visit comprised three main phases: familiarization, acquisition, and post-test (Figure 1). First, participants underwent a brief familiarization phase during which they were instructed to repeatedly and slowly perform the 8-element sequence until they accurately reproduced the sequence three consecutive times. This familiarization was intended to ensure that participants understood the instructions and explicitly memorized the sequence of finger movements. The actual acquisition phase consisted of physically performing 16 blocks of the 8-element motor sequence with their left (non-dominant) hand fingers. The first two blocks of this acquisition phase were used as a pre-test to evaluate baseline performance. Approximately five minutes after the end of the acquisition phase, all participants performed a post-test phase consisting of two blocks. This test was briefly preceded by a physical warm-up phase (i.e., slow-paced production of the sequence three consecutive times) to ensure that the correct sequence had been practiced. After the post-test (at 8:00 am), one-half of the participants had their left upper limb immobilized for 13 hours (IMMO group), while the other-half (CTRL group) had no restrictions on the use of their left limb. The immobilization kit consisted of an orthopedic splint immobilizing the wrist and 4 fingers (DONJOY brand, “Comfort Digit” model) and an immobilization sling (DONJOY brand, GCI model). To ensure that immobilization was carried out correctly and maintained throughout the day, an accelerometer (GENEactiv Original) was placed on each participant’s wrists, whether immobilized or not.

The second visit took place at 9:00 pm following the immobilization (or control) procedure at the Sleep and Vigilance Center of the Hôtel-Dieu hospital. After removing the immobilization for the IMMO group, all participants were administered another physical warm-up phase before performing a pre-night test consisting of two blocks of the 8-element sequence before sleep time. The EEG equipment was then set up for the second recording night. The next morning, at around 7:30 am and approximately 1 hour after awakening, participants carried out a physical warm-up phase before performing a post-night test consisting of two blocks of the 8-element sequence. Finally, the generalizability of the learned motor skill was assessed by conducting two transfer tests. An inter-manual transfer test was first administered to all participants to investigate skill generalizability toward the transfer from one limb to another. In this inter-manual transfer test, participants were required to perform two blocks on the original 8-element motor sequence with their unpracticed, dominant right-hand fingers (same keypresses). A transfer test with a new (unpracticed) 8-element sequence of stimuli (V-C-N-V-N-B-C-B) was then presented to differentiate sequence learning from generalized practice effects (67, 68). All participants were requested to perform this new motor sequence using their left (non-dominant) hand fingers. Each of these transfer tests was preceded by a brief familiarization phase during which participants were asked to perform the corresponding 8-element sequence (original or new) with the required hand slowly until they reproduced it correctly three times in a row.

### EEG-EMG data acquisition and pre-processing

#### EEG recordings

EEG was acquired using a 64-channel EEG cap (actiCAP snap BrainProducts Inc.) with slim-type electrodes (5kΩ safety resistor) suitable for sleep recordings. For reliable sleep stage scoring, electrooculography (EOG) and electromyography (EMG) recordings were added using bipolar Ag-AgCl electrodes. The EOG components were recorded by placing a pair of electrodes laterally to both eyes. EMG bipolar electrodes were placed over the chin. All EEG, EMG, and EOG data were recorded using two battery-powered 32-channel amplifiers and a 16-channel bipolar amplifier (respectively, BrainAmp and BrainAmp ExG, Brain Products Inc.). All signals were recorded at a 1-kHz sampling rate with a 100-nV resolution. Electrode-skin impedance was kept below 5 kΩ using Abralyt HiCl electrode paste to ensure stable recordings throughout all experimental phases.

EEG data were down-sampled to 250 Hz, bandpass filtered between 0.5 and 50 Hz to remove low-frequency drift and high-frequency noise, and re-referenced to the linked mastoids (i.e., TP9 and TP10). EOG and EMG data were respectively bandpass filtered between 0.3-35 Hz and 10-100 Hz.

### Statistical analysis

All subjects were included in statistical analyses (IMMO: n = 15; CTRL: n = 15), except as noted. Indeed, due to the absence of events of interest during some sleep EEG epochs, statistical power may be reduced for some specific analyses; this will be outlined where appropriate. The significant threshold was set at 0.05 for all analyses.

The sample size in each group was determined based on previous studies (9, 69, 70). Error measurements refer to the standard error of the mean (SEM) for non-circular data, and the circular standard deviation otherwise.

### Behavioral analysis

Response time (RT) was measured as the interval between two consecutive keypresses during each practice block. This performance index was described as more sensitive to the effects of sleep and the expression of relative performance gains (71, 72). Analyses regarding the number of accurately typed sequences are available in Supplementary Information (Figure S1). Also, since participants were asked to start over from the beginning of the sequence if they made any error during task production, RTs from error trials (i.e., erroneous key presses) were excluded from the analyses. To better reflect individual performance on the motor sequence task, mean RT performance was computed on accurately typed sequences (see (12, 73) for a similar procedure). Individual RTs were then averaged to obtain an overall estimation of the performance for each block. Motor skill consolidation was assessed by analyzing the overnight RT performance changes (in percentages) from the pre-night to the post-night test blocks. The inter-manual skill transfer was quantified by the RT performance changes (in percentages) between the post-night and the inter-manual transfer test blocks. Finally, the new-sequence skill transfer, or the capacity of transferring sequence knowledge to a new (unpracticed) sequence, was quantified by the RT performance changes (in percentages) between the post-night and the new-sequence transfer test blocks.

Mixed ANOVAs with the between-subject factor CONDITION (IMMO, CTRL) and the within-subject factor BLOCK were performed on the RT data to assess the difference between groups in the magnitude of overnight motor skill consolidation and generalization. Holm post-hoc comparisons were performed in case of significant effects or interaction.

### Actimetry analysis

To ensure that the upper limb was correctly immobilized, two accelerometers were placed on the participants’ wrists. The positions of both wrists were collected during the day between the acclimatization and experimental nights at a rate of 100 Hz. The velocity was computed across the day and averaged, giving us a mean velocity score for each upper limb. This score allows us to compare the degree of use of the immobilized left limb with the contralateral right limb for each participant in the IMMO group, as well as with the non-immobilized upper limbs of participants in the CTRL group. A mixed ANOVA was performed with the between-subject factor CONDITION (IMMO, CTRL) and the within-subject factor LATERALITY (Left, Right). Holm post-hoc comparisons were performed in case of significant effects or interaction.

### EEG analysis

#### Power spectrum density

The artifact-free EEG signal was sleep-stage scored according to AASM guidelines (74). Each 30-second epoch was visually scored as either NREM stages 1-3, REM, or wake. As described by Huber et al. (2006) (35), the immobilization procedure should induce local power decreases over sensorimotor regions mainly in the slow-oscillation frequency band (0.5-1.25 Hz) during the experimental night, at least in the first 20 minutes of sleep. Hence, for each participant and all electrodes, we restricted the EEG power spectrum density to the slow oscillation frequency band (0.5-1.25 Hz in steps of 0.25 Hz), computed using the Welch method (4-s Hamming window with 50% overlap between windows). The average power of the slow oscillation frequency band was obtained for all artifact-free NREM2 epochs of the first 20 minutes of sleep. A Student-t permutation test (1000 permutations) was performed to compare the power of the slow oscillation frequency band on each channel between the IMMO and CTRL groups. The statistical map was then corrected for multiple comparisons using the Benjamini-Hochberg procedure to control the false discovery rate (42) (Ntest = 63).

#### Spindle detection

Recent work has emphasized the greater contribution of NREM2 sleep spindle trains in motor memory consolidation following motor sequence learning (18). Hence, the detection of sleep spindle events was conducted using all artifact-free NREM2 sleep epochs over electrode C4 of the acclimatization and experimental night. This electrode was chosen based on the a priori hypothesis that immobilization of the left upper limb should affect the right sensorimotor cortex (35, 37). Discrete sleep spindle events (i.e., onset and offset) were automatically detected using a wavelet-based algorithm (see (18, 19) for further details). Spindles were detected at the C4 derivation by applying a dynamic thresholding algorithm (based on the mean absolute deviation of the power spectrum) to the extracted wavelet scale corresponding to the 11-16 Hz frequency range and a minimum window duration set at 300 ms (18, 19, 75) (see (8) for a review). Events were considered sleep spindles only if they lasted 0.3-2 seconds, occurred within the 11-16 Hz frequency range, and with onsets during NREM2 sleep periods. Visual inspections in random samples were done to ensure the correct detection of sleep spindles.

#### Spindle clustering

Sleep spindles may be split into two categories: clusters of two or more consecutive and electrode-specific spindle events interspaced by less than or equal to 6 seconds were categorized as trains, in comparison to those occurring in isolation (i.e., more than 6 seconds between two consecutive spindles detected on the same electrode) (8, 18). Hence, spindles belonging to trains were categorized as *grouped spindles*, and those occurring in isolation were categorized as *isolated spindles*. To evaluate the role of this temporal organization regarding sleep-related skill consolidation and generalizability, the proportion of grouped spindles defined by the total number of grouped spindles divided by the total number of spindles was computed. Hence, the greater the number of isolated spindles, the lower the proportion of grouped spindles. Pearson correlation analyses were performed to evaluate the relationship between the proportion of grouped spindles and the magnitude of overnight skill consolidation, inter-manual skill transfer, and new-sequence skill transfer. Fisher’s r to z transform was then performed to compare Pearson’s coefficients between groups.

#### Slow oscillation detection

Slow oscillation events were detected for each participant separately from sleep spindle events. The method used is similar to that described by Staresina et al. (2015) (30). First, the EEG data were bandpass filtered between 0.5-1.25 Hz (two-pass finite impulse response [FIR] bandpass filter). Only the artifact-free NREM2 and NREM3 epochs were considered. All zero-crossings were determined in the filtered signal, and SO candidates were identified as two successive positive-to-negative zero-crossings (i.e., down-states followed by up-states). SO events were retained if their duration was between 0.8 and 2 sec. Finally, the amplitude of the remaining SO events was determined. They were classified as SO if they met the amplitude criteria (≥ 75 % percentile of all SO amplitudes). Visual inspections in random samples were done to ensure the correct SO detection.

#### Phase-amplitude coupling

Sleep spindles and slow oscillations were classified as concomitant events if the onset of a sleep spindle occurred within the time interval of a slow oscillation (i.e., co-occurrence of SO and spindles). For each of these concomitant events, a Hilbert transform was applied to the extracted time window surrounding the SO onset (−1 to 2.5 seconds around the SO onset) to extract the instantaneous phase angle of the SO component (0.5-1.25 Hz) corresponding to the spindle power peak (11-16 Hz). For each participant, the average (preferred) phase θ between all concomitant events was given by the synchronization index (SI) (76).

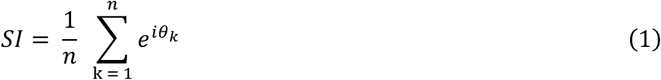

Where n is the number of concomitant events, θ_k_ is the SO phase value corresponding to the spindle’s power peak of the concomitant event k. The SI is a complex number whose angle corresponds to the preferred (average) phase of this synchronization, and its absolute value gives the average length of the resultant vector, from which the circular standard deviation is calculated. For illustration purposes, time-frequency maps in the sleep spindle frequency band were provided with the superimposed SO-filtered signal. For each extracted time window (−1 to 2.5 seconds around the SO onset), a time-frequency decomposition across the 10-20 Hz frequency band was performed using an eight-cycle Morlet wavelet (full width at half maximum; FWHM: 3 seconds for a central frequency of 1 Hz). Finally, the grand average TF map and SO-filtered signal were computed for each participant.

For each group, the distribution of the preferred phase was tested by the Rayleigh test. It is noteworthy that a non-uniform distribution of the preferred phase is an indicator of coupling (77). Watson-Williams tests were then used to compare the preferred phase of SO-spindle coupling between groups.

#### Temporal organization of sleep spindles

Two rhythms govern the temporal dynamics of sleep spindle activity: spindles tend to cluster on a low-frequency time scale of about 50 seconds (∼0.02 Hz infraslow rhythm), during which spindles iterate every 3-4 s (∼0.2-0.3 Hz mesoscale rhythm). *Infraslow rhythm.* The procedure to highlight the infraslow oscillation is similar to that described by Lecci et al. (2017) (22). To summarize, a continuous wavelet transform was performed on the artifact-free EEG data during the full night of sleep. The power time course was calculated in the 11-16 Hz range in steps of 0.2 Hz using a four-cycle Morlet wavelet (FWHM: 1.5 seconds for a central frequency of 1 Hz) and extracted for all artifact-free NREM2 sleep epochs. The average power in the spindle frequency band was calculated at each time point. A symmetric 4-second moving average was applied on the power time course to all consecutive 30-second NREM2 epochs, followed by its standardization, to reduce the temporal resolution and highlight the infraslow oscillation. A second continuous wavelet transform was performed on the power time course of the spindle frequency band, at a frequency resolution of 0.001 Hz in the 0.001-0.12 Hz range, and applied to all NREM2 epochs lasting more than 120 seconds (constituting a NREM2 period). Lastly, the spectral profile for each participant was computed by averaging the spectrum across all NREM2 periods with a 0.5-second time step, weighted by their duration. The final spectral profile per subject was obtained by normalizing it to its own mean. To gather the spectral profile peak for each participant and its corresponding frequency, a Gaussian fit (three terms) was performed, and the maximal value of the fitted curve was extracted. This maximal value, corresponding to the infraslow rhythm peak, was then compared between groups. *Mesoscale rhythm.* The same procedure was applied to study the mesoscale rhythm, except for the symmetric moving average which was not applied. After computation of the average power time course in the spindle frequency band, a second continuous wavelet transform was applied to all consecutive 30-second NREM2 epochs at a frequency resolution of 0.01 Hz in the 0.1-0.6 Hz range. Lastly, the spectral profile for each participant was computed by averaging the spectrum across all NREM2 periods with a 0.5-second time step, weighted by their duration. The final spectral profile was obtained by normalizing it to its own mean. A Gaussian fit (three terms) was performed, and the maximal value of the fitted curve was extracted. This maximal value, corresponding to the mesoscale rhythm peak, was then compared between groups.

Independent Student’s t-tests were performed between the IMMO and CTRL groups to compare the effects of the short-term upper-limb immobilization on the infralow and mesoscale rhythm peaks, respectively.

