## Supplemental Information for "Temporal clustering of sleep spindles and coupling with slow oscillations in the consolidation and generalization of motor memory"

**This PDF file includes:**

Tables S1 and S2

Figures S1 to S5

| | | CTRL | | IMMO | | Student's t-test | | |
|---|---|---|---|---|---|---|---|---|
| | | Mean | SEM | Mean | SEM | t | df | p |
| Whole night | NREM1 (min) | 45.2 | 3.5 | 32.5 | 2.6 | 2.87 | 28 | 0.008 |
| | NREM2 (min) | 186.6 | 5.9 | 172.0 | 7.5 | 1.52 | 28 | 0.14 |
| | NREM3 (min) | 99.5 | 5.6 | 87.2 | 5.7 | 1.49 | 28 | 0.15 |
| | REM (min) | 92.8 | 6.5 | 90.1 | 8.3 | 0.26 | 28 | 0.80 |
| | Total amount of spindles | 426.3 | 26.3 | 417.5 | 20.0 | 0.27 | 28 | 0.79 |
| | Total amount of grouped spindles | 184.7 | 16.9 | 182.7 | 12.3 | 0.11 | 28 | 0.92 |
| | Total amount of isolated spindles | 241.6 | 13.8 | 234.9 | 10.3 | 0.40 | 28 | 0.70 |
| | Proportion of grouped spindles (%) | 42.2 | 1.9 | 43.4 | 1.5 | -0.50 | 28 | 0.63 |
| First 20 minutes of NREM2 sleep | Total amount of spindles | 39.3 | 4.3 | 44.2 | 3.4 | -0.89 | 28 | 0.38 |
| | Total amount of grouped spindles | 20.9 | 2.8 | 25.7 | 3.1 | -0.80 | 27 | 0.43 |
| | Total amount of isolated spindles | 18.4 | 2.1 | 18.5 | 1.7 | -0.05 | 28 | 0.96 |
| | Proportion of grouped spindles (%) | 49.3 | 4.8 | 55.7 | 3.8 | -1.06 | 27 | 0.30 |

**Table S1. Sleep and spindle characteristics during the experimental night.** Means and standard errors of the means (SEM) are reported for the duration of NREM1, NREM2, NREM3, and REM sleep (in minutes), as well as the total amount of spindles, grouped spindles, isolated spindles, and the proportion of grouped spindles (in %) extracted at the C4 electrode from all NREM2 sleep epochs of the whole experimental night, for each group. Sleep spindle characteristics are also presented for the first 20 minutes of NREM2 sleep.

| | | CTRL | | IMMO | | Student's t-test | | |
|---|---|---|---|---|---|---|---|---|
| | | Mean | SEM | Mean | SEM | t | df | p |
| Whole night | NREM1 (min) | 44.5 | 3.4 | 41.9 | 5.4 | 0.42 | 28 | 0.68 |
| | NREM2 (min) | 173.4 | 11.0 | 175.7 | 12.2 | -0.14 | 28 | 0.89 |
| | NREM3 (min) | 79.4 | 5.3 | 73.3 | 7.2 | 0.68 | 28 | 0.51 |
| | REM (min) | 68.3 | 5.9 | 68.0 | 6.8 | 0.03 | 28 | 0.98 |
| | Total amount of spindles | 392.8 | 34.9 | 422.5 | 34.7 | -0.60 | 28 | 0.55 |
| | Total amount of grouped spindles | 167.6 | 18.7 | 180.3 | 15.3 | -0.53 | 28 | 0.60 |
| | Total amount of isolated spindles | 225.2 | 18.7 | 242.1 | 20.7 | -0.61 | 28 | 0.55 |
| | Proportion of grouped spindles (%) | 41.6 | 1.8 | 43.1 | 1.3 | -0.67 | 28 | 0.51 |
| First 20 minutes of NREM2 sleep | Total amount of spindles | 35.1 | 3.9 | 45.3 | 4.8 | -1.65 | 27 | 0.11 |
| | Total amount of grouped spindles | 18.3 | 2.9 | 25.1 | 3.5 | -1.52 | 26 | 0.22 |
| | Total amount of isolated spindles | 16.9 | 1.6 | 20.1 | 2.1 | -1.25 | 27 | 0.22 |
| | Proportion of grouped spindles (%) | 50.2 | 4.0 | 53.7 | 3.5 | -0.66 | 26 | 0.52 |

**Table S2. Sleep and spindle characteristics during the acclimatization night.** Means and standard errors of the means (SEM) are reported for the duration of NREM1, NREM2, NREM3, and REM sleep (in minutes), as well as the total amount of spindles, grouped spindles, isolated spindles, and the proportion of grouped spindles (in %) extracted at the C4 electrode from all NREM2 sleep epochs of the whole acclimatization night, for each group. Sleep spindle characteristics are also presented for the first 20 minutes of NREM2 sleep.

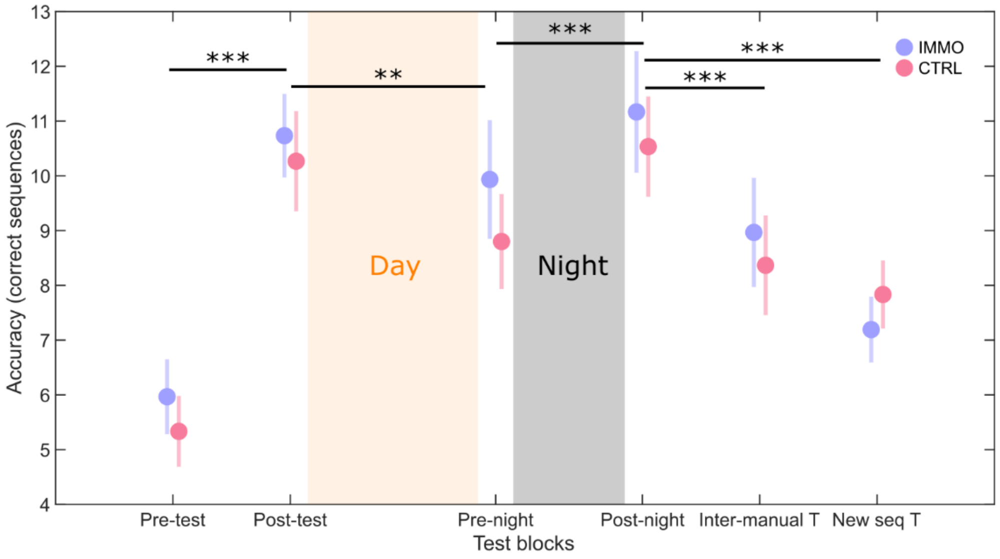


**Figure S1. Sensorimotor restriction did not affect motor sequence accuracy.** The colored circles represent the group average of the IMMO (purple) and CTRL (pink) groups, and the bar represents the standard error of the mean (n = 15 for each group and each test block, with the exception of the transfer test blocks with n = 13 for the IMMO group). Mixed ANOVAs using a CONDITION (IMMO, CTRL) x BLOCK factorial design, with repeated measures on the BLOCK factor, were performed to compare changes in performance over each interval. * $p < 0.05$, ** $p < 0.01$, *** $p < 0.001$. Inter-manual T: Inter-manual transfer; New seq T: new sequence transfer.

**Figure S2.** Given the a priori hypothesis that immobilization of the left upper-limb should affect the right sensorimotor cortex, mostly during the first 20 minutes of sleep, we assessed the difference in the power spectrum density between the IMMO and CTRL groups during the first 20 minutes of NREM2 sleep and the whole night at the C4 derivation. A significant SO power decrease was found for the first 20 minutes of NREM2 sleep (t(28) = -2.51, $p$ = 0.013, d = 0.92). However, the difference in the SO power between the two groups fades away when considering the whole night. To test this, a Student-t permutation test was performed between the groups on the average SO spectral power of NREM2 sleep for the whole night. The analysis failed to reveal a significant difference between the IMMO and CTRL groups (t(28) = -1.37, $p$ = 0.17, d = 0.50). Finally, two Student-t permutation tests for paired samples were performed to assess the evolution of the SO spectral power during NREM2 sleep between the first 20 minutes and the whole night for the IMMO and CTRL groups, respectively. Albeit not significant, our results revealed a trend towards significant SO power increases between the two time intervals (i.e., from the first 20 minutes to the whole night) in the IMMO group only (t(14) = -1.97, $p$ = 0.07, d = 0.51), not in the CTRL group (t(14) = -0.73, $p$ = 0.56, d = 0.19).

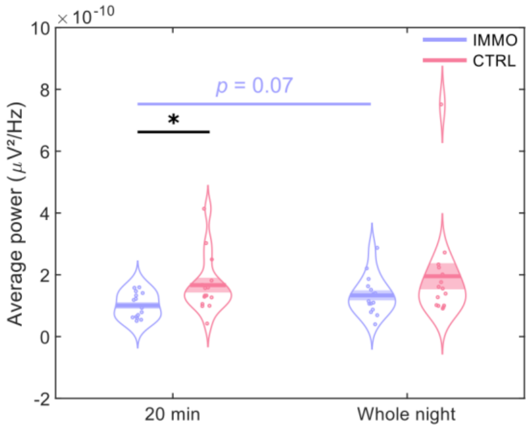


**Figure S2. Sensorimotor restriction affected slow oscillation activity in the first part of the night.** Comparison of the SO spectral power during NREM2 sleep between the first 20 minutes and the whole night for the IMMO (purple) and CTRL (pink) groups at scalp derivation C4. The curved lines indicate the distribution of data, the dark bars represent the mean of the distribution, and the lighter areas surrounding the mean represent the standard error of the means. Individual data points are displayed as colored circles. The star represents the p-value associated with the Student-t permutation test. * $p < 0.05$

**Figure S3.** Phase-amplitude coupling analyses were performed during the acclimatization night to assess whether the phase shift observed for the IMMO group during the experimental night is due to the immobilization procedure. These analyses were first performed on the SO-spindle concomitant events detected during the first 20 minutes of NREM2 sleep of the acclimatization night, extracted at scalp derivation C4. Participants without spindles coupled to slow oscillations during this period were removed from the current analysis (2 in the IMMO group, 3 in the CTRL group). The Rayleigh and the Watson-Williams tests were applied to assess the non-uniformity of the preferred coupling phases and to compare them between the IMMO and CTRL groups. Analyses indicated a non-uniform coupling in both IMMO ($\theta$ = -0.27 rad, Rayleigh Z = 8.31, $p < 0.001$) and CTRL groups ($\theta$ = -0.57 rad, Rayleigh Z = 8.21, $p < 0.001$). No significant difference in the preferred coupling phase was observed ($F(1,23) = 1.22$, $p = 0.28$), showing similar SO-spindle coupling for participants in both groups during the first 20 minutes of NREM2 sleep. Separate analyses of the SO-spindle coupling for grouped and isolated spindles were then performed on all artifact-free NREM2 epochs over the entire sleep recording of the acclimatization night.

First, for all NREM2 sleep spindles, a non-uniform SO-spindle coupling was found for participants in both IMMO ($\theta$ = -0.08 rad, Rayleigh Z = 14.1, $p < 0.001$) and CTRL groups ($\theta$ = -0.28 rad, Rayleigh Z = 13.3, $p < 0.001$). No significant group difference in the preferred phase was observed ($F(1,28) = 2.99$, $p = 0.095$).

Second, for all NREM2 grouped spindles, a non-uniform SO-spindle coupling was also found for participants in both IMMO ($\theta$ = -0.07 rad, Rayleigh Z = 12.7, $p < 0.001$) and CTRL groups ($\theta$ = -0.24 rad, Rayleigh Z = 12.3, $p < 0.001$). Again, no significant group difference in the preferred phase was observed ($F(1,28) = 1.10$, $p = 0.30$).

Third, for all NREM2 isolated spindles, a non-uniform SO-spindle coupling was also found for participants in both IMMO ($\theta$ = -0.09 rad, Rayleigh Z = 13.9, $p < 0.001$) and CTRL groups ($\theta$ = -0.31 rad, Rayleigh Z = 13.4, $p < 0.001$). No significant group difference in the preferred phase was observed ($F(1,28) = 3.70$, $p = 0.065$).

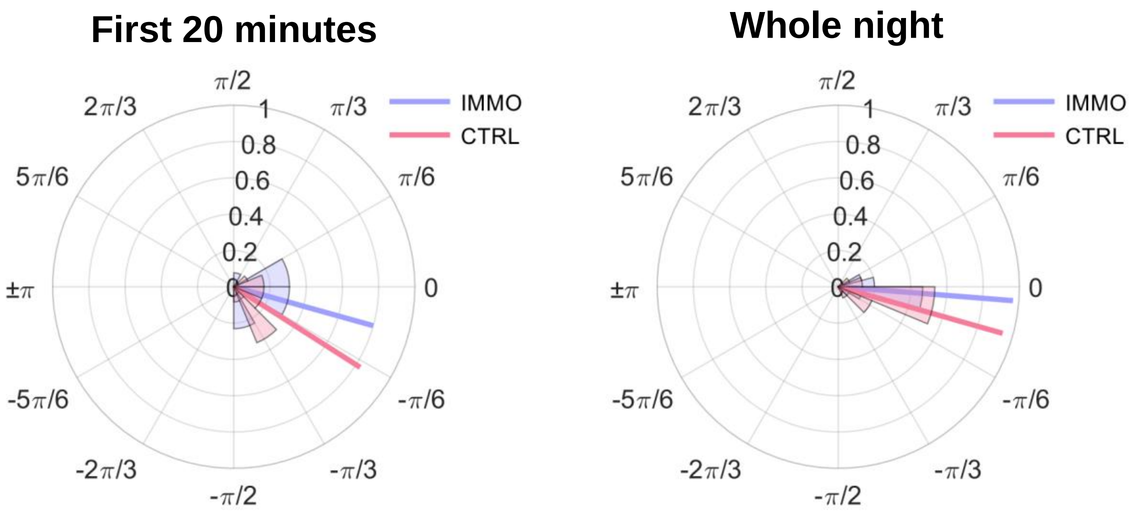


**Figure S3. Similar SO-spindle coupling during the acclimatization night for both groups.** Preferred phase in radians (and mean length of the resultant vector) of slow oscillations at the peak power of sleep spindles, detected at scalp derivation C4, for the IMMO (purple) and CTRL (pink) groups during NREM2 sleep when considering the first 20 minutes (left) and the whole night (right) of the acclimatization night. Probability density plots of phases (histograms) are also provided.

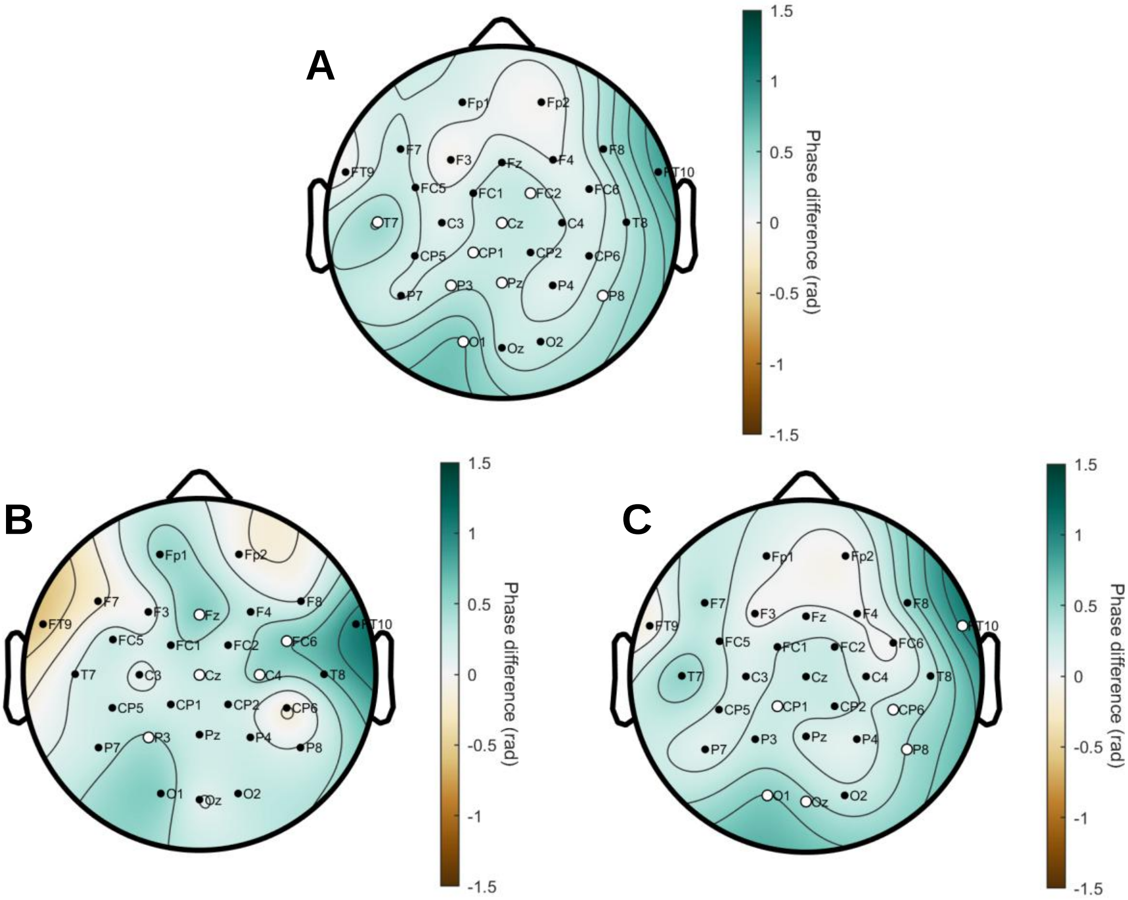


**Figure S4. Sensorimotor restriction induced local phase shift of the SO-spindle coupling.** The maps show the spatial variation of the SO-spindle coupling phase shift between the IMMO and CTRL groups (IMMO-CTRL contrast) during the experimental night for all NREM2 (**A**) sleep spindles (grouped and isolated altogether), (**B**) grouped spindles, and (**C**) isolated spindles. Green areas represent a delay in the coupling phase for the IMMO group, and conversely for brown areas. White dots highlight significant differences between the two groups in the preferred phase of the SO-spindle coupling after the Watson-Williams test.

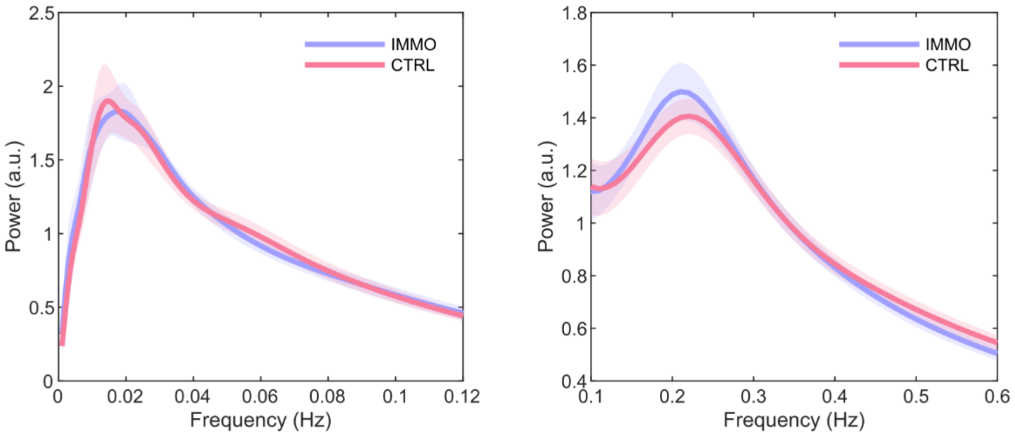


**Figure S5. Similar multi-scale fluctuations of spindle-band power during the acclimatization night for both groups.** Grand average of the spectral profile at the scalp derivation C4, ranging from 0.001 to 0.12 Hz (left panel) and 0.1 to 0.6 Hz (right panel) frequency range of the sleep spindle frequency band (11–16 Hz) power fluctuation during NREM2 sleep episodes across the whole acclimatization night. The peaks around 0.02 Hz and 0.2 Hz for both the IMMO (purple) and CTRL (pink) groups highlight the multi-scale periodicity of spindle-band power at an infraslow and mesoscale rhythms, respectively, before the immobilization procedure. No significant peak difference was found for these two rhythms between the IMMO and CTRL groups. The lighter areas surrounding the mean represent the standard error of the means.
